# Impacts of urbanization on the health of American Robins (*Turdus migratorus*) in Chicagoland

**DOI:** 10.1101/2024.11.24.625046

**Authors:** Anna Riccardi, Kimberly R. Douglass, Vera Soloview Jackson, Gavin K. Dehnert, Hannah Herbst, Felix Grewe, Morgan Walker, Seth Magle, Maureen H. Murray, Henry Adams, Cara E. Brook, Emily Cornelius Ruhs

**Affiliations:** Department of Ecology and Evolution, University of Chicago, Chicago, IL; Urban Wildlife Institute, Lincoln Park Zoo, Chicago, IL; Wisconsin Sea Grant; University of Wisconsin-Madison, Madison, WI; Grainger Bioinformatics Center, Field Museum of Natural History, Chicago, IL; Spatial Ecology and Epidemiology Research Laboratory, Department of Geography, University of Florida, Gainesville, FL

**Keywords:** health, immune function, parasite, stress, urbanization

## Abstract

Wild animals in urbanized environments face several unique challenges, including increased anthropogenic stressors, decreased natural food availability and quality, and increased pollutant exposure. While some work has shown that individual urbanization stressors can have negative impacts on aspects of wild bird physiology, other studies have demonstrated ambiguous or sometimes positive interactions. As such, the impact of multiple, coincident urban stressors on avian health still needs to be fully understood. Here, we addressed this knowledge gap by holistically measuring multiple physiological markers of American robin (*Turdus migratorius*) health across a gradient of urbanization throughout Chicagoland. We predicted that birds using highly urbanized habitats would experience higher heavy metal contamination, higher oxidative stress, lower body condition, higher avian malaria burden, and decreased measures of immune response compared to exurban birds in the Chicagoland area. Multiple linear models revealed that robins in more urbanized areas exhibited higher levels of heavy metal contamination and slightly elevated levels of associated physiological impairments compared to their counterparts in exurban sites. Additionally, noise and light pollution were significantly associated with oxidative stress and infection status, respectively, albeit in different directions. Overall, our findings underscore how the complex environmental changes that accompany urbanization can impact the health of urban bird populations.

## Introduction

Urbanization represents one of the greatest challenges that wild animals might encounter but can also provide ecological opportunities and refugia [1]. The expansion of urban landscapes forces wildlife to adapt to new environments or risk significant population losses [2–4]. In particular, some bird species have documented declining abundance, reproductive ability, and survival in urban areas [5]; however, suboptimal natural conditions can sometimes provide increased anthropogenic foods and lessen energetic burden [1]. With over half of the world’s human population living in urban environments, and this number projected to increase [3], it is important to consider how the complex and multifaceted presence of human activity could impact wildlife health.

While many factors could influence the physiology of birds in urbanized environments, some major challenges urban birds face include low-quality diets [2] and increased light and noise pollution [6–8]. In urban environments, birds are known to forage in refuse, consume secondary anthropogenic food sources, or be directly fed human food [9–11], which can result in lower antioxidant micronutrients [2,12] leading to increased oxidative damage [13]. Invertebrate food sources are also less available in less green areas [12] and more polluted environments [14]. In addition to altered resources, chronic exposure to elevated noise and light pollution can increase stress hormone levels and disrupt the balance between pro-oxidants and antioxidants, resulting in higher oxidative stress in some urban-dwelling birds [15,16]. Therefore, urbanization may negatively impact overall body condition and immune function [17]. Passeriform birds found in urban settings can have higher parasite burdens ([18,19] though some studies have identified the opposite [20]), lower constitutive immunity [21], and lower body condition [21]. Continued stress, low levels of antioxidants, and impaired immunity are just a few of the challenges that birds living in heavily urbanized environments face, potentially leading to altered physiological parameters [22,23].

Birds residing in anthropogenic environments also experience exposure to environmental contaminants, particularly heavy metals. Several studies have documented higher levels of heavy metal pollutants in urban areas than in rural environments [2,24]. Common sources of heavy metal contamination in cities include industrial activities, vehicle emissions, improper waste disposal, and the weathering of infrastructure containing metals like lead and zinc [25,26]. Heavy metals can accumulate in urban soils, vegetation, and invertebrate prey items, thus entering the food web and bioaccumulating in higher trophic-level species (e.g., birds). Moreover, some heavy metals (e.g., lead, mercury and cadmium) can negatively impact avian physiology by inducing oxidative stress, neurotoxicity, endocrine disruption, and impaired immune function [26,27]. Therefore, while urbanization stressors (e.g., light, noise pollution and environmental contaminants) represent significant challenges for birds inhabiting urbanized areas, we need to learn more about how these factors combined influence bird health.

Herein, our goal was to understand the impacts of urbanization on the overall health and physiology of a resident bird species, the American robin (*Turdus migratorus*), in the Chicagoland region. We sampled birds from seven sites representing an urban-to-exurban gradient in the Chicagoland area to investigate the relationship between proxies of health status and exposure to various anthropogenic stressors such as artificial light, noise, and heavy metal contamination. We predicted that highly urbanized habitats would have greater levels of anthropogenic stressors, associated with a resident bird population with higher oxidative stress and impaired immunity than exurban birds in the Chicagoland area. We characterized the level of anthropogenic stressors that could threaten birds by quantifying the amount of light pollution, noise pollution, and heavy metals exposure across an urban-rural gradient in Chicagoland and evaluated associations with estimated defenses (antioxidant capacity), innate immune function, stress (H:L ratio), and avian malaria parasite infection in individual American robins. By enhancing our understanding of the physiology of American robins in urban habitats, we can identify anthropogenic stressors that may be targeted to reduce health-related stress in urban birds. This understanding can also guide future urban development to minimize potential negative physiological impacts on wildlife.

## Materials and Methods

### Determination of urban-rural gradient

To create a composite urban intensity metric for Chicagoland, we first followed previously reported methods [28] to apply a principal component analysis (PCA) to canopy cover (%) (CMAP, 2018) [29], impervious cover (%) (CMAP, 2018), and housing density (derived from U.S. Census, 2010) within 1000m of the mist-netting sites. These variables were extracted in R v 4.0.4 (R Development Core Team, 2021) via the uwinspatialtools package [30]. We retained the first PCA component (PC1), which explained 88.8% of the variation in these data, as a measure of urban intensity (UI). The loadings of PC1 were canopy (-0.60), impervious (0.56), and housing density (-0.56). Because this approach resulted in a negative metric, to make it more intuitive as a measure of urban intensity, we multiplied all values by -1. Thus, this metric represents a gradient of urban intensity (hereafter, “urban intensity”) with lower values corresponding to less urban regions dominated by greenspace and higher values corresponding to the most urban regions dominated by built environments with dense housing.

### Capture and sampling

We surveyed seven sites (Wolf Road Woods, Blackwell Forest Preserve, Beck Lake, Miller Meadow, Ottawa Trail Woods, Jackson Park, Columbus Park) across the rural to urban landscape in the Chicago metropolitan area for American robins. Robins were captured following a similar method as described before [31]. All mist netting was conducted in 2020 and 2021 from mid-August through mid-October. Mist nets were checked every 15-30 minutes and non-target species were extracted and immediately released without collecting samples. American robins were extracted and placed in individual cloth bags to reduce handling stress. The bird was weighed (±0.1 g), tarsus length and wing chord were measured (±0.1 mm), and banded with a U.S. Geological Survey aluminum band. Each captured bird was sexed and aged, a blood sample (see blood collection methods), body feathers (∼25 body feathers in the breast region) were collected, and then the bird was released at the site of capture. Site-related information and sample sizes can be found in Table 1. Permits were obtained to mist-net and band birds under U.S. Geological Survey Federal Bird Banding Permit #09924.

**Table 1.** Site-related summary of anthropogenic stressors and contamination levels. Noise is the estimated proportion exposed to 50 to 60 dB. Light is represented by “night sky brightness” - Zenith radiance (mcd/m^2^). Housing density is housing units per kilometer squared Canopy cover is percent of landcover. Impervious surface is percent of landcover.

| Site name | Site abbreviation | County | Latitude | Longitude | Sample size<br>Total (males;<br>females) | Noise | Light | Housing<br>density | Canopy<br>cover | Impervious<br>surface | Urban<br>intensity |
| --- | --- | --- | --- | --- | --- | --- | --- | --- | --- | --- | --- |
| Wolf Road Woods | WRW | Cook | 41.70453 | -87.89525 | 11 (4; 7) | 5.8 | 3.8 | 0.32 | 0.79 | 0.01 | 2.44 |
| Blackwell Forest<br>Preserve | BLW | DuPage | 41.82822 | -88.17656 | 2 (1; 1) | 4.6 | 8.7 | 154.77 | 0.47 | 0.1 | 1.11 |
| Beck Lake | BKL | Cook | 42.07051 | -87.87218 | 4 (3; 1) | 38.2 | 10.4 | 582.14 | 0.46 | 0.17 | 0.55 |
| Ottawa Trail Woods | OTW | Cook | 41.81527 | -87.80465 | 21 (10; 11) | 96.9 | 26.9 | 820.02 | 0.47 | 0.29 | 0.02 |
| Miller Meadow | MMW | Cook | 41.86229 | -87.82883 | 5 (3; 2) | 13 | 34.7 | 377.69 | 0.29 | 0.27 | -0.2 |
| Jackson Park | JKP | Cook | 41.7863 | -87.57977 | 12 (7; 5) | 69.2 | 59.5 | 1655.01 | 0.25 | 0.24 | -1.32 |
| Columbus Park | CLP | Cook | 41.87546 | -87.76732 | 19 (11; 8) | 20.5 | 80.4 | 2660.69 | 0.22 | 0.5 | -2.59 |

### Blood collection, transport, and processing

For each bird, a blood sample was taken (≤1% of the bird’s body mass) via jugular puncture using a sterile syringe with an attached needle (26 gauge). Blood smears were created [32] for white blood cell counts (see WBC count methods). Blood was kept cold and centrifuged within 6 hours of collection to separate the serum from the red blood cell pellets. Both blood cell pellets and serum were then stored in a -80°C freezer until further analyses were performed.

### Body condition calculation

Body condition in avian species has been calculated in numerous ways [33], and the “best” measure is species dependent. Here, we calculated the linear correlation between mass (in grams) and (1) tarsus length (mm) and (2) wing chord (mm), while controlling for age class (Fig S1). While both relationships were at least marginally significant between mass and measurement, we ultimately opted to use tarsus length in downstream analyses to avoid any confounding measures due to feather wear (Fig S1). To derive an expression for body condition, we used the linear model form (mass ∼ total tarsus length) separately by age group (adult, young), and calculated the predicted mass based on structural size. We then subtracted the predicted mass from the actual mass to get a measure of mass:tarsus length residuals; negative values corresponded to lower body condition scores and positive values to higher body condition scores.

### Molecular sexing

Birds were sexed molecularly via PCR using the CHD gene and the CHD1F 5’-TATCGTCAGTTTCCTTTTCAGGT-3’ and CHD1R 5’-CCTTTT ATTGATCCATCAAGCCT-3’ primer pair [34]. Briefly, reactions occurred in 27μl volumes with 8.5μl H20, 12.5μl GoTaq Green master mix (Promega), 1.0μl forward primer (100μM), 1.0μl reverse primer (100μM), and 4.0μl DNA template per reaction. Samples were run at 95° (5min), [94°C (30s)48°C (30s)72°C (60s)] for 28 cycles, 72°C (5min). Males were identified by observing one band for the Z-chromosome, and females were identified by observing two bands, one band for the Z-chromosome and one band for the W-chromosome.

### White Blood Cell Counts

Blood smears were air dried, fixed with 100% methanol, and stained with Wright-Giemsa stain [32]. Cells were viewed with the Leica DM500 microscope under 100x magnification with oil immersion to determine the leukocyte profile [35]. Lymphocytes, heterophils, eosinophils, monocytes, basophils, and thrombocytes were counted. The total number of white blood cells (WBCs) were reported out of 10,000 red blood cells. If unable to distinguish between heterophils and eosinophils, cells were counted as granulocytes. The total WBC count and heterophil to lymphocyte (H:L) ratio were used for data analysis as a proxy for stress [36]. If there were multiple slides per bird, the average count between the slides was used for statistical analysis.

### Avian Malaria Parasitism

Avian malaria is a mosquito-borne infection caused by the blood parasite, *Plasmodium relictum* [37]. *P. relictum* is endemic to North America, including the Chicagoland area, and can cause high rates of mortality or chronic morbidity in birds [37]. For each blood smear, the number of red blood cells infected with blood parasites were counted out of 10,000 red blood cells. Three common blood parasites of birds were screened: *haemoproteus*, *plasmodium*, and *leucocytozoon*. Each bird was screened for blood parasites using three primer sets with nested PCR protocols. PCR1 targeted *haemoproteus* and *plasmodium* parasites [38], PCR2 targeted *haemoproteus*, *plasmodium*, and *leucocytozoon* [39], whereas PCR3 only targeted *leucocytozoon* [40]. For PCR1, we based our protocol on [38] using first-round primers HAEMNF and HAEMNR2 and second-round primers HAEMF and HAEMR2. For PCR2, we based our protocol on [39] using first-round primers HAEMNF1 and HAEMNR3 and second-round primers HAEMFL and HAEMR2L (Supplementary Table S1). For PCR3, we based our protocol on [40] using first-round primers DW2 and DW4 and second-round primers LeucoF and LeucoR. This combination of primer sets was implemented to reduce the high rate of false-negative detections commonly occurring under only one nested protocol (*per communication*). We used the same master mix as in the molecular sexing protocol above; however, in nested PCRs, 4.0µl of PCR template 1 from the first-round was used as the template for the PCR of the second round. For specific primer sequences and thermocycler profiles for each PCR protocol see Table S1. Samples which displayed a band at the ∼520bp region for any of the three PCR runs were submitted for Sanger sequencing at the University of Chicago DNA sequencing core (summary information can be found in Table S5).

### Circulating IgY antibodies

Serum samples were used to assess circulating baseline antibody (IgY) levels, an important part of innate immunity which can shed insight on later memory-mediate antibody protection, using an enzyme-linked immunosorbent assay (ELISA) [23]. In short, we coated a 96-well plate with 1µl serum diluted in 99µl coating buffer - in duplicate - and incubated the plate overnight (∼16 hours) at 37℃. After the incubation, the coating solution was removed, 200µl of block buffer was added to each well and then incubated at room temperature for 30 minutes. The plate was then washed four times with ELISA wash buffer. Then we added 50µl of goat anti-bird IgG (Bethyl A140-110P) diluted in 1:1000 blocking buffer and incubated at 37℃ for 1 hour. The plate was again washed four times. 100µl of ABTS substrate was added to each well and incubated at room temperature in the dark for five minutes. 100µl of 1% SDS was added across the plate and read with an Epoch spectrophotometer at λ = 405nm.

### Antioxidant Capacity

An oxy-adsorbent assay was used to measure circulating antioxidant capacity by testing the functional capacity of components in the plasma to neutralize an introduced oxidant. We used the Diacron OXY-Adsorbent test (Diacron International # MC435, Grosseto, Italy) to measure the capacity of plasma to neutralize the pro-oxidant hypochlorous acid (HClO). In short, 5µl of each plasma sample was diluted 1:100 in 495 µl of distilled water. In a 96-well plate, 200 µl of the R1 reagent (tittered HClO solution) was aliquoted, followed by 5µl of the diluted plasma, 5µl distilled water for a control, or 5µl of calibrator serum in duplicate. After incubation at 37°C for 10 minutes, 2.5 µl of the R2 reagent (N,N-diethyl-p-phenylenediamine solubilized in a chromogenic mixture) was added to each well. The plate was then read with a spectrophotometer at λ = 540nm. We then calculated the concentration of HClO that was neutralized as an indication of the impairment degree of the antioxidant barrier of the sample. The calibrator serum has a known antioxidant capacity of 350 mmol/l of HClO neutralized; the absorbance read of the calibrator and control distilled water are used in the following equation, as directed by the manufacturer.

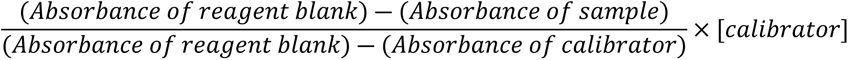

Measurements were expressed as mmol/l of HClO neutralized.

### Analysis of sound and light pollution

We employed the National Transportation Noise Exposure Map (NTNM) to assess the impact of noise pollution on robin populations across various sampling sites in the Chicagoland area [41,42]. This dataset integrates noise levels stemming from aviation, roadway, and rail traffic, offering a detailed landscape of urban noise pollution. The NTNM uses the 24-hour equivalent A-weighted sound level (LAeq) at a 30 m x 30 m resolution as its metric. We focused on noise levels of LAeq between 50 to 60 dB to assess the impact on robin populations. For night sky brightness (e.g., light pollution), the tiff and tpk files contain the entire floating-point dataset from the 2015 World Atlas of Artificial Night Sky Brightness. The data represent the artificial component of the night sky brightness (natural sky brightness is not included).

### Heavy Metal Analysis

For each bird and site, breast feathers were analyzed at the USGS Mercury Research Laboratory (USGS-MRL, Madison, WI) to quantify mercury concentrations. Prior to analyses, all samples were lyophilized and then homogenized by ball mill. For methylmercury analysis, approximately 15mg of dried tissue were weighed into extraction vials, to which 4mL of 4.5 M HNO3 was added and then sealed with Teflon-lined closures. The sample extracts were heated at 55°C for 12 hours and then analyzed by a Brooks Rand Merx-M automated methylmercury analysis instrument [43]. For total mercury analysis, approximately 10mg of tissue was weighed into combustion boats and analyzed directly using a Nippon MA-3000 Direct Combustion Analyzer following EPA method 7473 [44].

Quality assurance and control measures were employed throughout sample analysis at the USGS-MRL, including standard reference material (SRM) and triplicate sample analyses. For tissue analyses, methylmercury recovery of SRM IAEA 407 (fish homogenate) averaged 105.5% (n = 3, 99.9% - 112.0%) and sample precision assessment by triplicate analyses was 3.1% and 7.0% (relative standard deviation). Total mercury SRM recovery of IAEA 407 averaged 103.1% (n=34, 97.5% - 108.2%) and sample precision assessment by triplicate analyses averaged 1.3% relative standard deviation (n = 8, 0.2% - 3.5%). For sediment analysis, total mercury recovery of sediment SRM IAEA 456 averaged 97.8% (n=4, 95.1% - 99.7%) and was not assessed for precision. All sample results exceeded lab method detection limits for both methylmercury and total mercury analysis in tissues (1.3 ng/g and 3.8 ng/g, respectively) and for total mercury analyses in sediments (1 ng/g).

For all other heavy metal analyses, 100 uL of hydrogen peroxide was added to the prepared sample described above. Prepared samples were allowed to dissolve for 7 days. 2mL of the prepared sample was added to 6.5mL of 2% nitric acid and run through a 0.45um Agilent filter. Concentrations of 26 heavy metal elements (Ag, Al, As, B, Ba, Be, Ca, Cd, Co, Cr, Cu, Fe, K, Mg, Mn, Mo, Na, Ni, Pb, Sb, Se, Si, Ti, Tl, V, and Zn) were analyzed in all samples. Analysis of the feather samples was performed using an inductively coupled plasma optical emission spectrometer (ICP-OES). The ICP-OES used was a Spectro Arcos (SPECTRO, Germany) with Torch type of Flared end EOP Torch 2.5 mm. Operating optimal parameters were: RF generator (1400 W), Argon was plasma, auxiliary and nebulizer gas. Plasma gas flow, auxiliary gas flow, and nebulizer gas flow were 14.5, 0.9 and 0.85 (L/min), respectively. After that, sample uptake time, rinse time, initial stabilization time was 240 total, delay time and time between replicate analysis were zero. The measurement replicate was three-time, the frequency of the RF generator was 27.12 MHz (resonance frequency). Type of detector solid state and spray chamber were CCD and Cyclonic, Modified Lichte, respectively. Sample delivery pump type was four-channel, software controlled; peristaltic pump enables exact sample flows. Prewash pump speed (rpm) was 60 (for 15 s), 30 (for 30 s) and prewash time was 45 s at the end sample injection pump speed was 30 rpm. Results are reported in the appropriate units for the analysis and metal type.

### Statistical Analysis

#### Contaminants and health metrics across urbanization intensity

A series of generalized linear models (GLMs) with bird toxicant contamination level as the response variable and urban intensity scores (e.g., PC1*-1) as a fixed factor were run to determine significant differences in various contamination levels across an urbanization gradient. We completed the same approach for each of the bird health metrics across an urbanization gradient. We included one additional model to understand if parasite infection covaried with urbanization intensity. Due to the number of GLMs being completed for the health metrics, we followed up with a Benjamini-Hochberg correction to control the expected proportion of Type 1 errors among the rejected hypotheses. We did not perform this correction for the level of contaminants, as we consider individual contaminants to be independent of one another, unlike the health metrics.

#### Contaminants across sampling sites

Because we only found a marginal effect of urban intensity on the levels of heavy metal contamination, we then wanted to understand if there were site-level differences (factors) across contaminants to be able to explore the spatial differences of those sites. Therefore, a series of Analysis of Variances (ANOVAs) with toxicant name as the response variable and site ID as a fixed factor were run to determine significant differences in various contamination levels across our sampling sites with a TukeyHSD post-hoc test run on all subsequently significant ANOVAs.

#### Health metrics across sampling sites

Because we did not find an impact of urbanization intensity on health metrics, we then ran a series of ANOVAs with assay type as a response variable and site ID as a fixed factor for ten health metrics (Body condition, Oxidative stress, Total WBC count, IgY, H:L ratio, and percentages of all individual WBC types: Lymphocyte, Heterophil, Monocyte, Basophils, and parasite prevalence across the urbanization gradient with a TukeyHSD post-hoc test run on all subsequently significant ANOVAs. All p-values were adjusted using a holms-Bonferroni approach, as we did not expect health metrics to be independent.

#### Influence of heavy metal contamination and urban stressors on health metrics

We found no association between urbanization intensity and bird health, but subsequently attempted to identify whether specific factors associated with different sites were associated with different metrics of bird health. Thus, to assess the direct relationship between urban stressors (e.g., heavy metal contamination, light pollution, and noise pollution) and metrics of robin body condition and health, we ran a series of linear models on select metrics (body condition, antioxidant capacity, total WBC count, IgY and H:L ratio) with the physiological metric as the response variable, the concentrations of each metal with suspected impacts on animal health (Al, As, Cd, Cr, Cu, Mn, Pb, Zn, Hg ug/g) [45–48], light at night (mcd/m^2^), noise (dB), sex, and age as fixed factors. For each of these models, we considered the distribution family to be gaussian (e.g. normally distributed), except for total WBC (poisson) and H:L ratio (inverse gaussian). We also included a final logistic regression model to understand what urban stressors, sex, and age influenced parasite infection status (0/1 infected) with a binomial distribution.

## Results

### Site selection from least to most urbanized

By using the PCA analysis, we categorized the sites from rural to urban in the following order, based on the urban intensity scores : (7) Wolf Road Woods (6) Blackwell Forest Preserve, (5) Beck Lake, (3) Ottawa Trail Woods, (4) Miller Meadow, (2) Jackson Park, (1) Columbus Park (Fig 1; Table 1). In subsequent figures, the sampling site is ordered accordingly from least urbanized to most urbanized.

**Fig 1.**
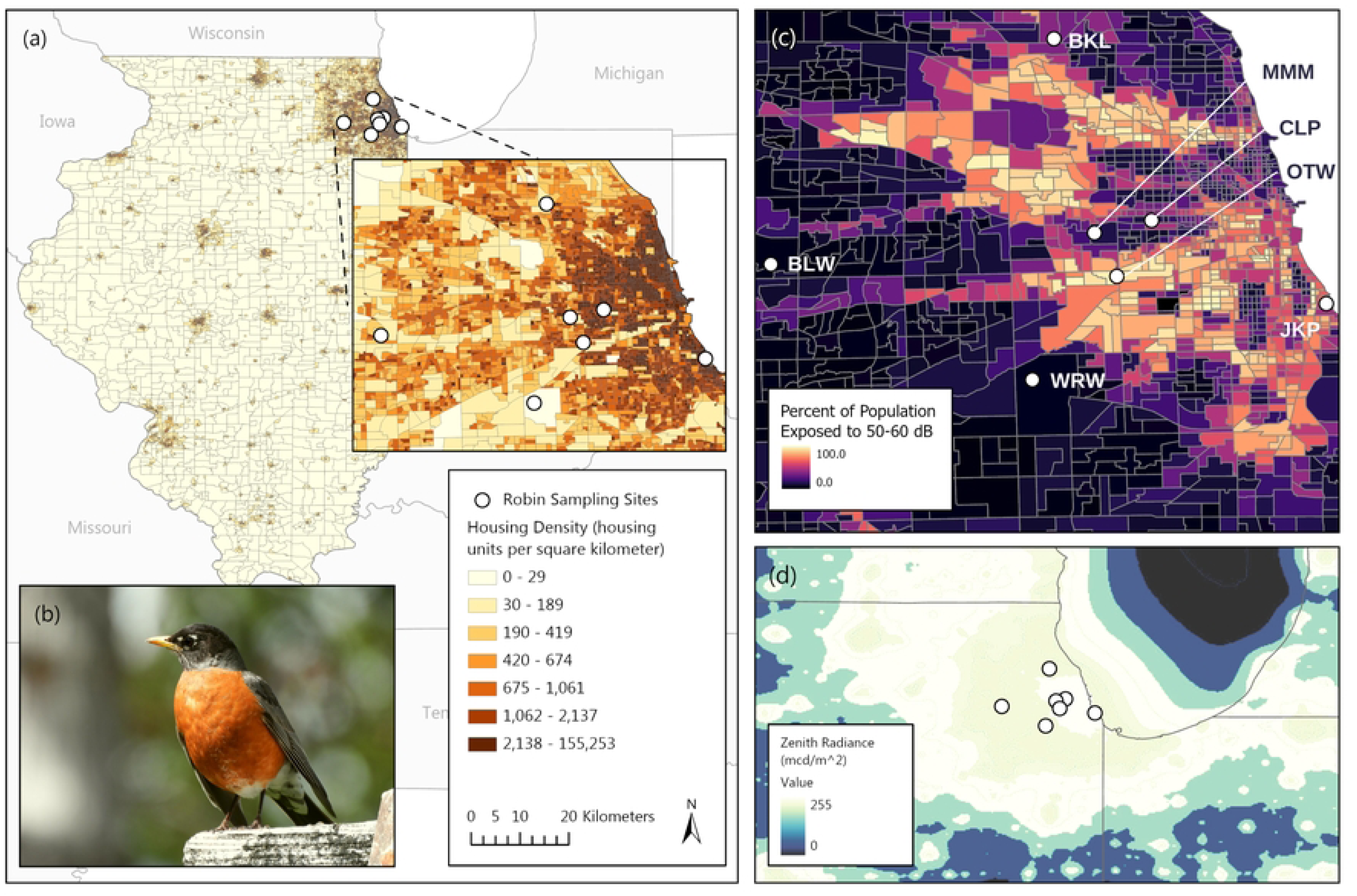
Sites of American robin sampling by the Lincoln Park Zoo and Urban Wildlife Institute. (a) Housing density of Illinois, with the inset of Chicagoland with points representing sampling sites. (b) An American Robin in an urban setting. (c) Noise pollution of Chicagoland as determined by the percent of the population exposed to over 50-60dB. (d) Light pollution of Chicagoland as determined by the Zenith radiance (mcd/m^2^).

### Contaminants and health metrics across urbanization intensity

Of the twenty-seven heavy metals assessed, only Thallium (Tl) showed significant increases in concentration with increases in urbanization (Table S2). None of the health metrics measured were associated with urbanization intensity (Table S3).

### Contaminants across sampling sites

Twenty-seven different heavy metal concentrations (ug/g) were analyzed in the American robins’ feathers (Fig S2). The majority of heavy metals had consistent levels across the seven sites; however, four heavy metals had significant differences across sites: Barium (Ba), Mercury (Hg), Sodium (Na), and Zinc (Zn) (Fig 2; Table S4). Barium concentrations in robin feathers were significantly higher in Beck Lake compared to Blackwell Forest Preserve, Wolf Road Woods, Ottawa Trail Woods, Jackson Park, and Columbus Park (Table 1). Sodium concentrations in robin feathers were significantly higher in Beck Lake compared to Jackson Park. Mercury concentrations in robin feathers were significantly higher in Jackson Park compared to Wolf Roads Woods, Blackwell Forest Preserve, Ottawa Trail Woods, Miller Meadow, and Columbus Park. Zinc concentrations in robin feathers were significantly higher in Beck Lake compared to all other sites. Some heavy metals showed trends that correlated with the predicted urban gradient (e.g., arsenic, iron) while others went against the predicted trend (e.g., sodium, potassium; Fig S2).

**Fig 2.**
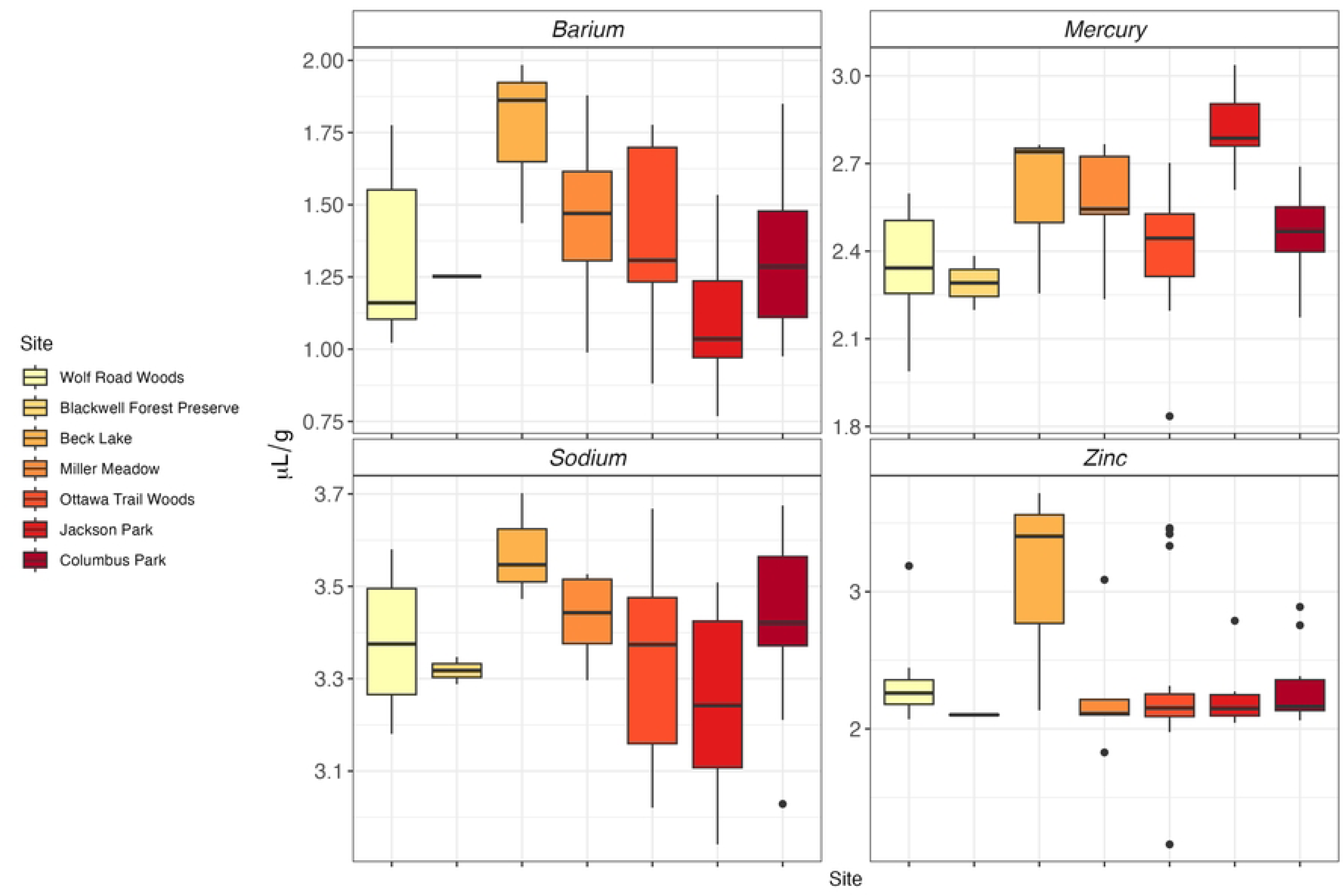
Heavy metal contamination across Chicagoland sites. Contamination levels were significantly different for Barium, Mercury, Sodium, and Zinc; but not for the other 18 metals analyzed (Fig S1).

### Health metrics across sampling sites

Ten health metrics were analyzed to assess robin health (Fig S3). After statistical correction, only one was significant: the percentage of heterophils (Fig 3a; Table S5). The percentage of heterophils were significantly lower in Columbus Park than in Ottawa Trail Woods and Jackson Park.

**Fig 3.**
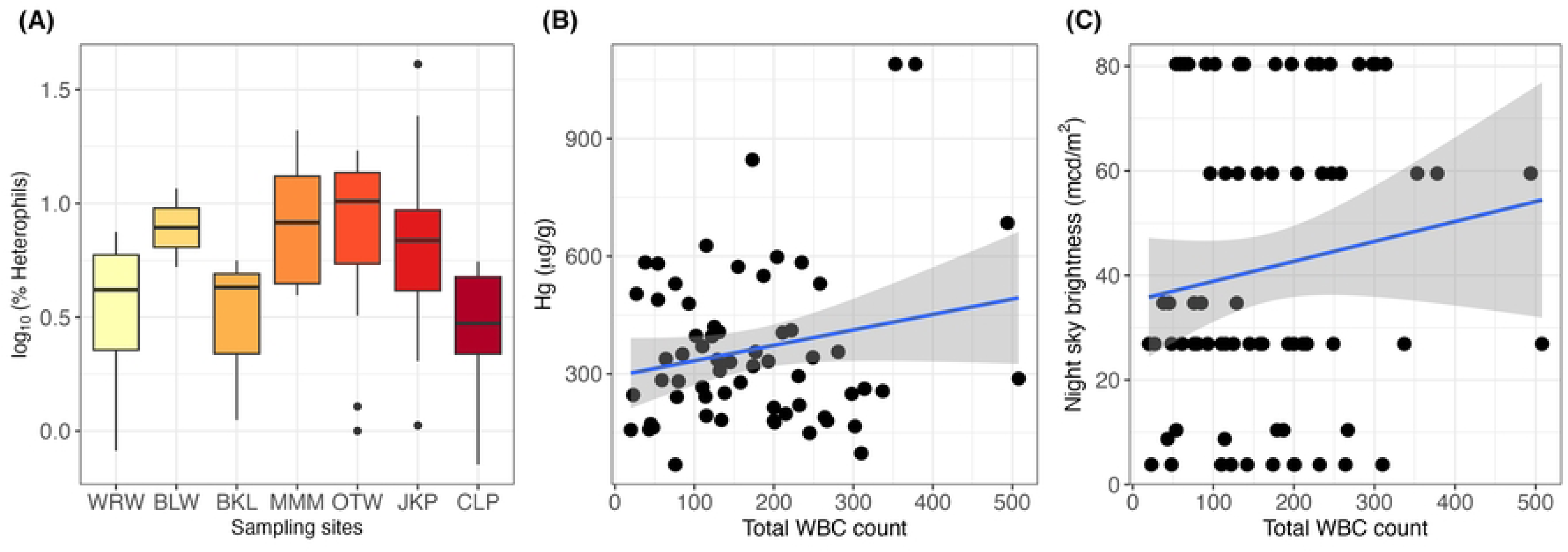
Physiological condition of American robins sampled across Chicagoland sites. **(A)** Physiological condition was significantly different among sites for the percentage of heterophils; but not for the other nine physiological metrics assessed. Sites are organized from least to most urbanized. Total WBC counts increased with both **(B)** increasing mercury (Hg) concentration and **(C)** increasing night sky brightness.

### Influence of urbanization stressors on health metrics

We wanted to understand whether specific factors (e.g. light, noise, contamination) of the sites were associated with different metrics of bird health. This was important because no strong trends of changes in health metrics occurred across the urbanization gradient (above). However, when we ran linear models to determine the explanatory urbanization stressors of various physiological metrics, only the total WBC count appeared to be influenced by levels of select contaminants, light pollution, and noise pollution (Table S6). The direction of the relationship between total WBC count and concentrations of heavy metals were highly dependent on the exact pairing: total WBC count increased with increasing cadmium, manganese, lead, zinc, and mercury concentrations (Fig 3b), but declined with increasing aluminum, arsenic, chromium, and copper concentrations. The clearest patterns were with extraneous measures of urbanization such that increasing total WBC counts were associated with greater light pollution (Fig 3c). Additionally, H:L ratios varied by sex of the bird (e.g., with ratios being higher in females).

Parasite prevalence varied greatly across sampling site and was as follows: Columbus Park=63.16%, Jackson Park= 83.33%, Miller Meadow=80%, Ottawa Trail Woods=85.71%, Beck Lake=75%, Blackwell Forest Preserve=100%, and Wolf Road Woods=100% (Fig 4a). It should be noted that sample sizes were quite low for Miller Meadow (n=5) and Blackwell Forest Preserve (n=2) and should be interpreted with caution. From the initial linear model, birds in more intense urban areas were more likely to be infected with avian malaria (β=0.5347 ± 0.2254, z=2.372, p=0.0177; Fig 4b). Sanger sequencing results revealed that almost all parasite species were of *Plasmodium spp.* (Table S7).

**Fig 4.**
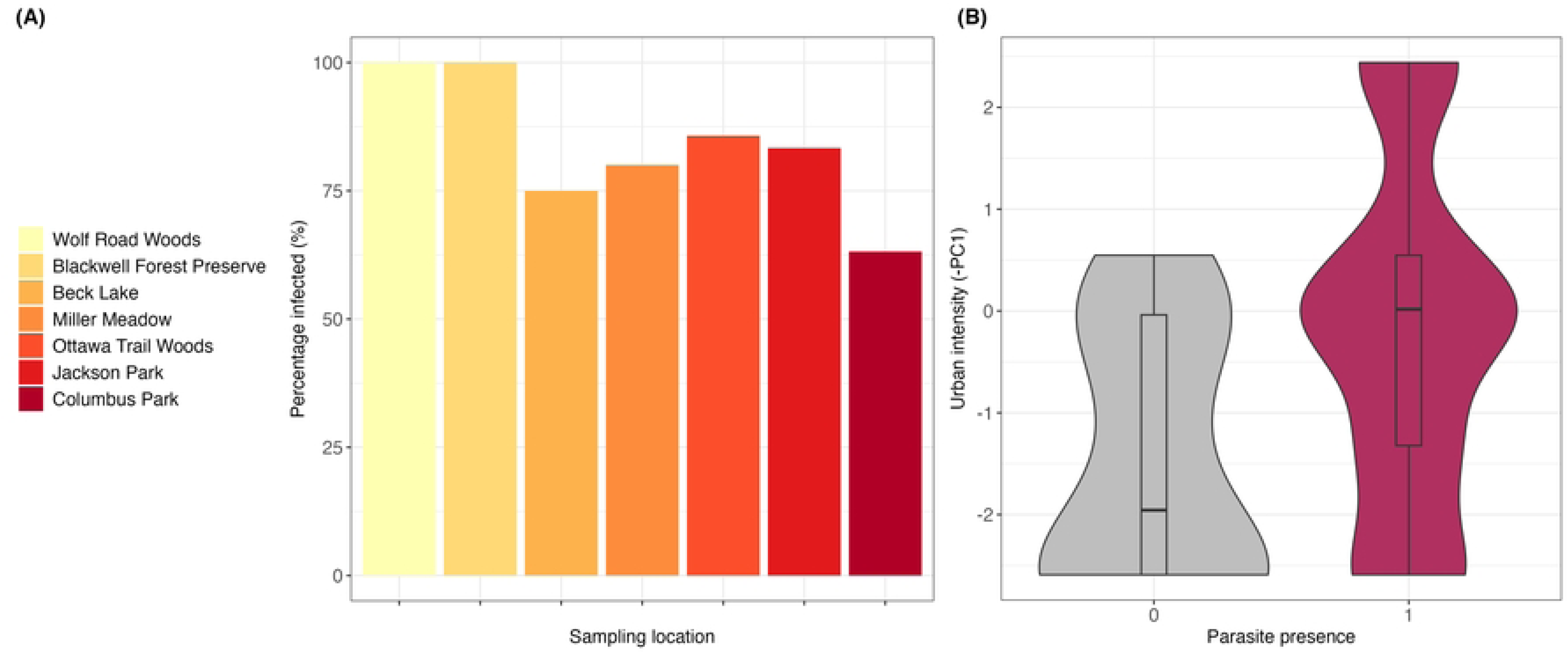
Influence of urban stressors on parasite infection and prevalence. **(A)** The prevalence of avian malaria infection in sampled American robins was high throughout Chicago land and demonstrated no significant association across the urbanization gradient (with details provided in the main text); despite this, **(B)** generalized linear models demonstrated that parasite infection in birds was associated with a greater urbanization intensity.

## Discussion

Our study offers novel insights into the physiological challenges facing urban wildlife by examining relationships between anthropogenic stressors associated with urbanization (e.g., noise pollution, light pollution, contaminant exposure) and markers of health, oxidative status, and parasite prevalence in American robins across a rural-urban gradient in the Chicago metropolitan region. While we did not find many significant associations between the degree of urbanization and physiological condition, our results revealed complex spatially-heterogeneous patterns. Site-specific trends emerged with robins in more urbanized locations generally exhibiting greater burdens of mercury contamination, coupled with altered immune profiles and other indicators of compromised health, such as percentage of heterophils. These findings highlight how urbanization’s environmental impacts could lead to complex physiological alterations and potentially increased morbidity.

In general, the levels of heavy metals were comparable to other studies on bird feathers in large urban areas [49,50]. Barium, mercury, sodium, and zinc were the only heavy metals that differed by site of capture. Interestingly, excluding the two least urbanized sites (Blackwell Forest Preserve and Wolf Road Woods), which experienced similar levels of barium, there was a clear downward trend in barium levels as the degree of urbanization increased. Barium is a naturally occurring element; however, a significant portion of the barium in the environment comes from coal combustion, primarily found in suburban and less densely populated areas [51]. Barium is also used in a variety of industrial applications, including the production of automotive paint, fireworks, and certain industrial activities (e.g., the production of glass, ceramics, and some chemicals) [52]. Sodium followed a similar trend, with the level of contamination generally decreasing as the degree of urbanization increased. The application of sodium-based fertilizers and pesticides in agricultural settings could also lead to elevated sodium levels in nearby waterways and ecosystems [53]. Overall, the barium and sodium data indicate that American robins in more rural, exurban areas of Chicago are experiencing greater exposure to these heavy metal pollutants than their urban counterparts.

In contrast to the patterns observed for barium and sodium, mercury contamination levels increased with the level of urbanization on the site level. In the Chicagoland area (which has a long history of industrial processes), there are recorded levels of “legacy sources” of mercury; historical industrial activities and waste disposal practices have led to the accumulation of mercury-contaminated soils and sediments [25]. Furthermore, the Clean Air Mercury Rule permanently capped mercury emissions from coal-fired power plants in 2005, steadily reducing the amount of mercury released into the environment [50]; thus, this result is likely driven by the historic impact of mercury on the site-level. Additionally, thallium increased with increasing urban intensity. Similarly to mercury, thallium is also released into the environment through industrial sources (e.g., coal burning) and because thallium has higher water solubility compared to other heavy metals, it can be more bioavailable and bioaccumulate [54]. Last, we found that zinc contamination was relatively low at most sites; however, there was a stark exception at Beck Lake, where feather samples showed high concentrations of zinc. This localized hotspot suggests the presence of a point source for zinc pollution in the area surrounding Beck Lake - potentially due to sunscreen use at the recreational site [55] or proximity to a regional airport [56]- which contrasted with the minimal zinc exposure observed for robins at the other sites sampled throughout the Chicago metropolitan area. The differences in the patterns of these heavy metal results underscore that microscale bioaccumulation patterns of metals can be complex.

In the present study, we found measurable correlations between the degree of urbanization and a few key physiological health indicators in American robins across the Chicagoland area. Heterophils trended towards a decrease with urbanization when compared by site, but not urbanization intensity (e.g., -PC1), however, this effect was not significant and may be due to the limited urbanization gradient captured by our seven sites. Future studies may better examine this relationship by increasing the number of sites studied and capturing a larger gradient of urbanization. We also observed a correlation between many heavy metals and total WBC count - an association previously observed [49,50]. Specifically, we observed an increase in total WBC count with higher mercury contamination and greater light pollution (night sky brightness). Levels of heavy metals have been previously shown to impact specific leukocyte compositions in birds, especially heterophils and lymphocytes [57–59]. Kidney mercury levels, specifically, were associated with increased percentage of heterophils in two species of water birds (*Aechmophorus occidentalis* and *A. clarkii*) [58].The observed relationships between urbanization and these physiological markers are likely driven by the complex interplay of various anthropogenic stressors that increase as habitats become more developed, including chemical pollutants, noise pollution, light pollution [60]. Future studies should aim to tease apart these potentially counter-acting sources of pollution and the overall contribution to the relationship with urbanization.

Finally, when we examined whether factors of urbanization influenced avian malaria presence, we found that birds from more urbanized sites, and thus greater light pollution, experienced higher parasite prevalence than those birds sampled from rural areas. This same pattern was reflected for *Leucocytozoons* but not for *Haemoproteus* or *Plasmodium*, in Eurasian blackbirds (*Turdus merula*) and house sparrows (*Passer domesticus*) [58,61]. Interestingly, a global investigation of *Plasmodium vivax* (one of four human malaria parasites) reported the rate (Malaria Atlas Project) was much lower in urban areas compared to rural areas [62]. Interestingly, studies of human malaria have shown higher prevalence of infection in less urban environments, reflecting the habitat preferences of Anopheles spp. mosquito vectors. Avian Plasmodium is vectored instead by Culex spp. mosquitoes, which have more urban habitat preferences. Continued monitoring across a larger range of urbanization of these types of sublethal physiological responses can provide valuable insights to guide conservation and management efforts aimed at mitigating the impacts of urbanization on avian communities.

## Conclusion

Ultimately, birds across the Chicago metropolitan area, even in nominally “natural” exurban sites, are exhibiting slight physiological imprints of anthropogenic stress and contamination, but the patterns are highly complex [63]. The American robins we sampled displayed evidence of high parasite prevalence, some altered immunity, and accumulated burdens of anthropogenic contaminants, even at localities touted as ecological refugia. Our findings underscore the need for continued concerted regional efforts to mitigate sources of anthropogenic stress - from tightening industrial emissions controls to remediating contaminated sites to reducing light, noise, and anthropogenic food pollution, especially in major metropolitan regions like Chicagoland. Comprehensive interventions are necessary to curtail negative impacts to bird survival across human-altered landscapes [28].

## Acknowledgements

We would like to thank members of the Brook and Dehnert labs for feedback on previous versions of this manuscript. We would also like to acknowledge our funding sources: A.R. received a Quad Scholars grant from the University of Chicago, V.S.J. received a Dean’s Fund research grant and a BSCD Ecology and Evolution Fellowship, E.C.R was funded by a Grainger Bioinformatics Foundation fellowship.

---

**Fig S1. Mass residuals against morphometric structures.** Tarsus length was marginally significantly correlated to mass for adults (p=0.11) and young (p=0.09). Wing chord was marginally significantly correlated to mass for adults (p=0.09) and significant for young (p=0.015).

**Fig S2. Heavy metal contaminant concentration (µg/g) across sites.**

**Fig S3. Physiological metrics in American robins sampled across sites.**

**Table S1.**
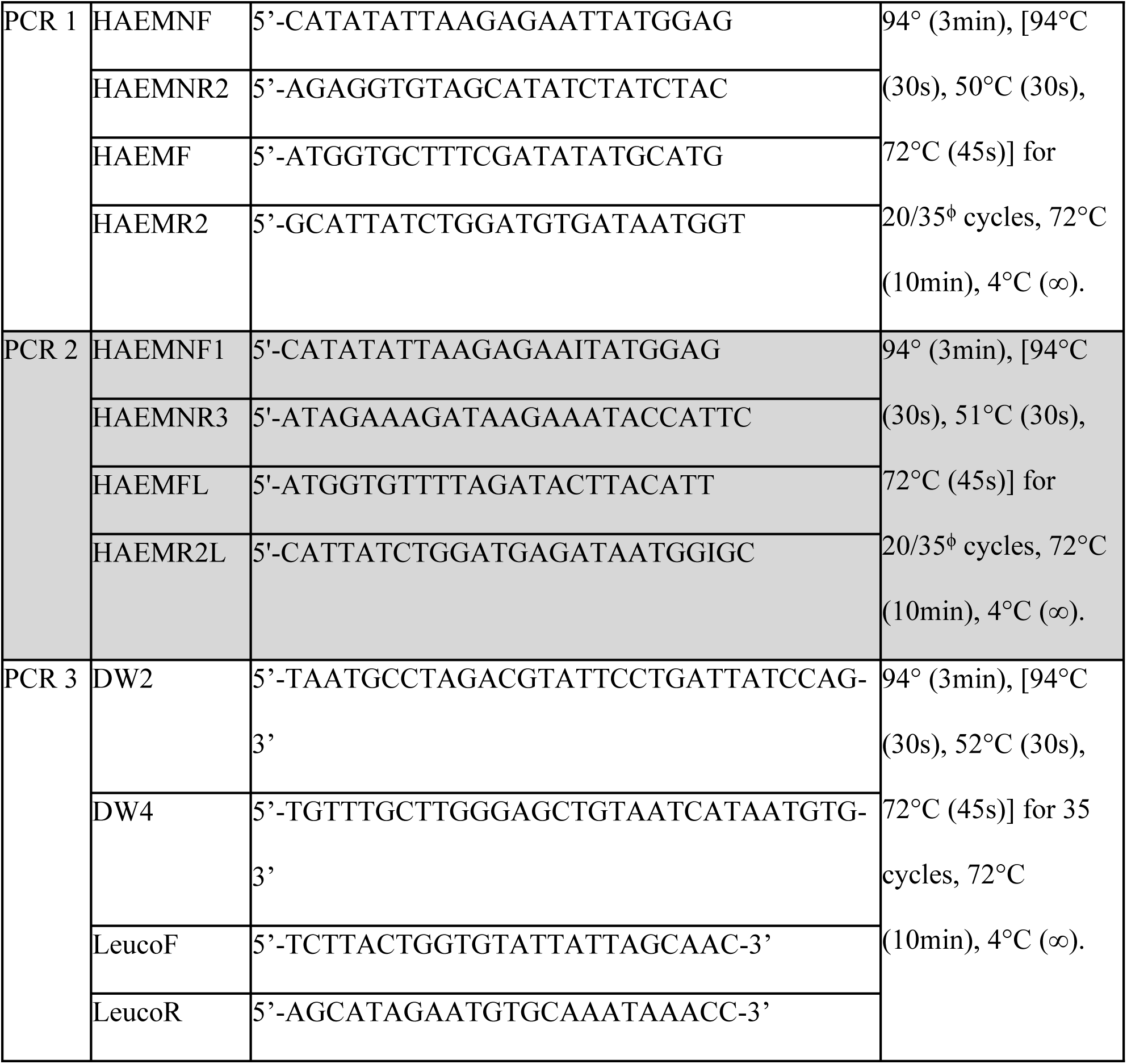
Protocol primer sets and thermocycler profiles for each PCR protocol. ^ϕ^The number of cycles is the only aspect of the thermocycle profile that changed between first and second rounds.

**Table S2.** Results of the generalized linear models for the effect of urbanization on heavy metal levels. Because we considered heavy metals to be independent responses, we did not apply a correction to the models. All heavy metals highlighted in bold are considered significant at P<0.05.

| Toxicant name | Estimate | Std. error | t value | P value |
| --- | --- | --- | --- | --- |
| Aluminum | -3.926 | 10.116 | -0.388 | 0.699 |
| Antimony | -1.052 | 1.173 | -0.897 | 0.373 |
| Arsenic | -0.3724 | 0.4406 | -0.845 | 0.401 |
| Barium | 0.990 | 1.667 | 0.594 | 0.555 |
| Boron | -19.17 | 60.14 | -0.319 | 0.751 |
| Calcium | -81.48 | 52.36 | -1.556 | 0.125 |
| Cadmium | -0.1624 | 1.0605 | -0.153 | 0.879 |
| Chromium | -5.710 | 4.2 | -1.36 | 0.178 |
| Copper | -0.4652 | 0.3996 | -1.164 | 0.248 |
| Iron | -82.73 | 43.14 | -1.918 | 0.0594 |
| Lead | 0.2455 | 0.6760 | 0.363 | 0.718 |
| Magnesium | -9.617 | 18.844 | -0.51 | 0.611 |
| Manganese | 0.3788 | 0.6445 | 0.588 | 0.559 |
| Mercury | -30.27 | 15.87 | -1.908 | 0.0611 |
| Potassium | 13.39 | 20.32 | 0.659 | 0.512 |
| Selenium | -0.1565 | 0.4519 | -0.346 | 0.73 |
| Silver | 7.823 | 4.065 | 1.925 | 0.0584 |
| Sodium | -26.95 | 71.24 | -0.378 | 0.706 |
| Silicon | -16.36 | 16.43 | -0.996 | 0.323 |
| **Thallium** | **1.0494** | **0.4351** | **2.412** | **0.0186** |
| Titanium | -0.5293 | 0.3897 | -1.358 | 0.179 |
| Zinc | 68.60 | 64.63 | 1.061 | 0.292 |

**Table S3.** Results of the generalized linear models for the effect of urbanization on health metrics. P-values are adjusted for multiple comparisons using a Benjamini-Hochberg correction. Urbanization intensity was not a predictor of any health metric we measured.

| Assay name | Estimate | Std. error | t-value | P value | Adjusted P value |
| --- | --- | --- | --- | --- | --- |
| Body condition | 0.4533 | 0.3243 | 1.398 | 0.166 | 0.9393135 |
| Oxidative stress | -0.7384 | 4.1867 | -0.176 | 0.861 | 0.9393135 |
| IgY | 0.00099 | 0.0061 | 0.163 | 0.871 | 0.9393135 |
| Heterophil:Lymphocyte | 0.00399 | 0.01828 | 0.218 | 0.828 | 0.9393135 |
| Total WBC | -6.905 | 7.276 | -0.949 | 0.346 | 0.9393135 |
| Lymphocyte (%) | -1.353 | 1.989 | -0.68 | 0.499 | 0.9393135 |
| Heterophil (%) | 0.3104 | 0.4604 | 0.674 | 0.502 | 0.9393135 |
| Eosinophil (%) | -0.03683 | 0.4221 | -0.087 | 0.931 | 0.9393135 |
| Monocyte (%) | 0.04569 | 0.59810 | 0.076 | 0.939 | 0.9393135 |
| Basophil (%) | 0.0075 | 0.052 | 0.144 | 0.886 | 0.9393135 |

**Table S4.** Results of the ANOVAs identifying contamination differences across sites. Results are reported for the effect of the site. Heavy metals are all in µg/g. Significant differences are indicated by asterisks and bolded. Results of the post-hoc analysis and specific difference between sites are listed in the main text.

| Toxicant name | Sum Sq | Mean Sq | F value | P value |
| --- | --- | --- | --- | --- |
| Aluminum | 210607 | 35101 | 2.024 | 0.0754 |
| Antimony | 2891 | 481.9 | 2.053 | 0.0715 |
| Arsenic | 255.9 | 42.65 | 1.202 | 0.317 |
| **Barium** | **88784** | **1479.0** | **3.501** | **0.00473** |
| Boron | 5289263 | 881544 | 1.364 | .243 |
| Calcium | 2033153 | 338859 | 0.683 | 0.664 |
| Cadmium | 1350 | 225 | 1.095 | 0.375 |
| Chromium | 16995 | 2833 | 0.837 | 0.546 |
| Copper | 281.1 | 46.85 | 1.651 | 0.148 |
| Iron | 2047173 | 341196 | 0.94 | 0.473 |
| Lead | 411 | 68.43 | 0.798 | 0.575 |
| Magnesium | 729238 | 121540 | 2.106 | 0.0766 |
| Manganese | 553 | 92.11 | 1.222 | 0.307 |
| **Mercury** | **1534360** | **255727** | **12.66** | **5.774e-9** |
| Potassium | 906375 | 151062 | 2.176 | 0.0569 |
| Selenium | 151.4 | 25.23 | 0.65 | 0.69 |
| Silver | 30204 | 5034 | 1.612 | 0.158 |
| **Sodium** | **11604544** | **1934091** | **2.299** | **0.0453** |
| Silicon | 292247 | 480708 | 0.963 | 0.458 |
| Thallium | 262 | 43.66 | 1.171 | 0.333 |
| Titanium | 287.6 | 47.94 | 1.783 | 0.117 |
| **Zinc** | **16921773** | **2820296** | **4.8** | **0.0004** |

**Table S5.** Results of the ANOVAs identifying differences in health metrics across sites. Results are reported for the effect of the site. Significant differences are bolded. Adjusted p-value represents the holm’s-bonferroni correction for multiple comparisons. Results of the post-hoc analysis and specific difference between sites are listed in the main text.

| Assay name | Sum Sq | Mean Sq | F value | P value | Adjusted P value |
| --- | --- | --- | --- | --- | --- |
| Body condition | 2.8232e+02 | 4.7053e+01 | 2.442 | 0.0338 | 0.3038 |
| Oxidative stress | 3.6227e+04 | 6.0378e+03 | 1.915 | .0915 | 0.4495 |
| IgY | 7.871e-03 | 1.3118e-03 | 0.1714 | 0.9836 | 1.0 |
| Heterophil:Lymphocyte | 7.7161e-01 | 1.286e-01 | 2.1019 | 0.0642 | 0.4495 |
| Total WBC | 1.1959e+05 | 1.9932e+04 | 2.0209 | 0.0747 | 0.4495 |
| Lymphocyte (%) | 1.0022e+04 | 1.6704e+03 | 2.333 | 0.0416 | 0.3326 |
| **Heterophil (%)** | **7.3871e+02** | **1.2312e+02** | **3.4795** | 0.0047 | **0.047** |
| Eosinophil (%) | 1.0826e+02 | 1.8043e+01 | 0.487 | 0.8158 | 1.0 |
| Monocyte (%) | 2.7021e+02 | 4.5035e+01 | 0.6118 | 0.7201 | 1.0 |
| Basophil (%) | 5.7119e+00 | 9.5198e-01 | 1.9519 | 0.0853 | 0.4495 |

**Table S6.** Results of the models identifying the relationship between physiological metrics, toxicant concentration, and urban stressors. Results are reported for the significant effects only; thus, only models with significant predictors are shown.

| Physiological metric | Fixed effect | Estimate | Std. error | z value | P value | Adjusted P value |
| --- | --- | --- | --- | --- | --- | --- |
| Heterophil:Lymphocyte | Sex | 2.695e-01 | 7.007e-02 | 3.846 | 0.00078 | 0.0217 |
| Total WBC | Aluminum (ugg) | -5.73 e-4 | 1.445e-04 | -3.969 | <0.0001 | 8.89 e-5 |
|  | Arsenic (ugg) | -3.14 e-2 | 2.517e-03 | -12.479 | <0.0001 | 2.59 e-35 |
|  | Cadmium (ugg) | 1.73 e-2 | 1.268e-03 | 13.644 | <0.0001 | 9.21 e-42 |
|  | Chromium (ugg) | -3.56 e-3 | 2.503e-04 | -14.207 | <0.0001 | 6.6 e-45 |
|  | Copper (ugg) | -1.4 e-2 | 2.713e-03 | -5.228 | <0.0001 | 2.28 e-7 |
|  | Manganese (ugg) | 2.37 e-2 | 1.554e-03 | 15.253 | <0.0001 | 2.5 e-51 |
|  | Lead (ugg) | 9.13 e-3 | 1.533e-03 | 5.957 | <0.0001 | 3.74 e--9 |
|  | Zinc (ugg) | 1.93 e-4 | 2.239e-05 | 8.612 | <0.0001 | 1.44 e-17 |
|  | Mercury (ugg) | 5.19 e-4 | 4.893e-05 | 10.614 | <0.0001 | 5.86 e-26 |
|  | Light | 6.26 e-3 | 4.590e-04 | 13.64 | <0.0001 | 9.21 e-42 |
| Parasite infection | Urban Intensity | 0.5347 | 0.2254 | 2.372 | 0.0177 |  |

**Table S7.**
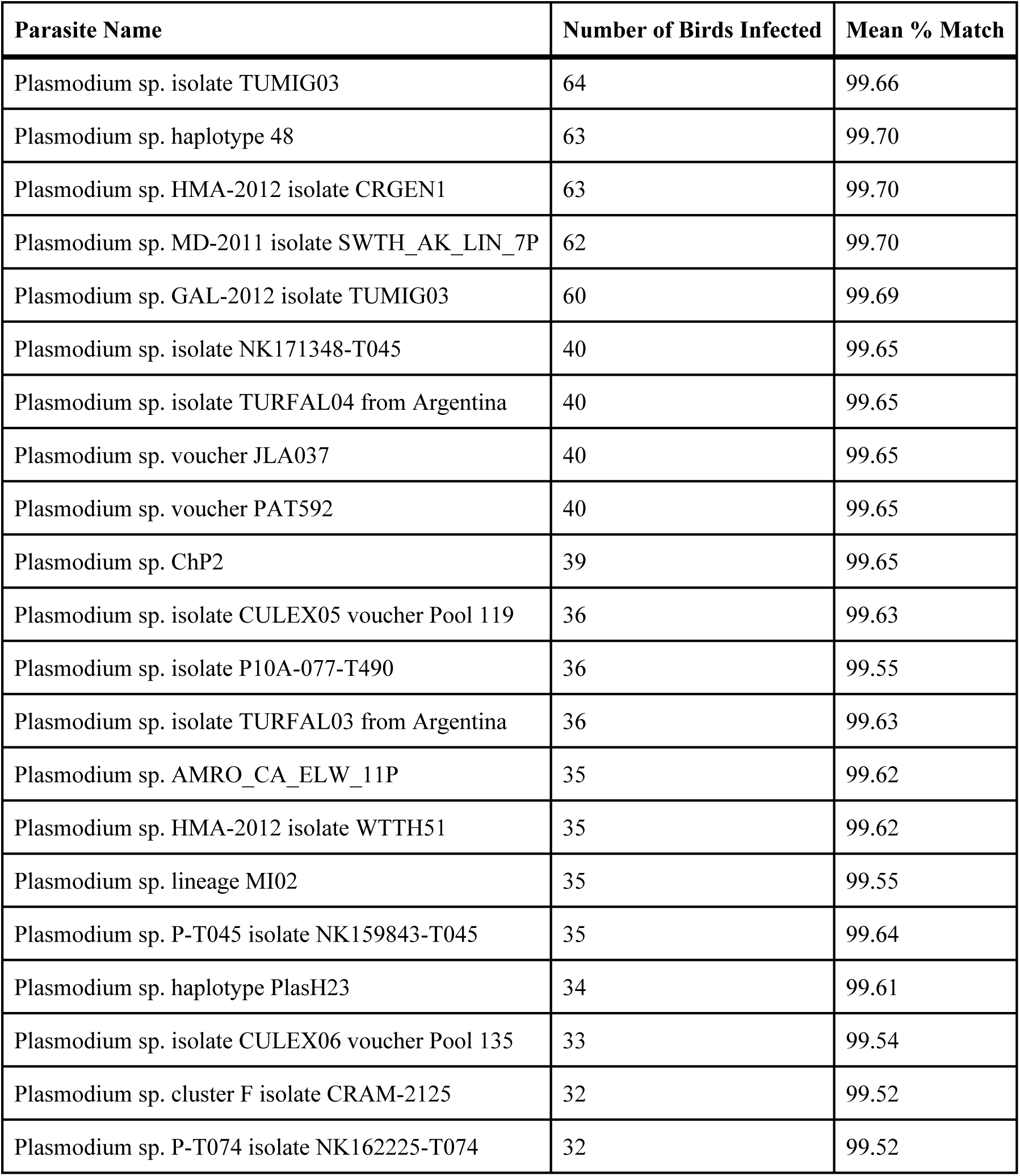
Results of the sanger sequencing of PCR avian malaria positives.

